# Automating an insect biodiversity metric using distributed optical sensors: an evaluation across Kansas, USA cropping systems

**DOI:** 10.1101/2023.08.15.553397

**Authors:** Klas Rydhmer, James O. Eckberg, Jonathan G. Lundgren, Samuel Jansson, Laurence Still, John E. Quinn, Ralph Washington, Jesper Lemmich, Thomas Nikolajsen, Nikolaj Sheller, Alex M. Michels, Michael M. Bredeson, Steven T. Rosenzweig, Emily N. Bick

## Abstract

Global ecosystems and food supply depend on insect biodiversity for key functions such as pollination and decomposition. High-resolution, accurate data on invertebrate populations and communities across scales are critical for informing conservation efforts. However, conventional data collection methodologies for invertebrates are expensive, labor intensive, and require substantial taxonomic expertise, limiting researchers, practitioners, and policymakers. Novel optical techniques show promise for automating such data collection across scales as they operate unsupervised in remote areas. In this work, optical insect sensors were deployed in 20 agricultural fields in Kansas, USA. Measurements were compared to conventional assessments of insect diversity from sweep nets and Malaise traps. Species richness was estimated on optical insect data by applying a clustering algorithm to the optical insect sensor’s signal features of wing-beat frequency and body-to-wing ratio. Species richness correlated more strongly between the optical richness estimate and each of the conventional methods than between the two conventional methods, suggesting sensors can be a reliable indicator of invertebrate richness. Shannon- and Simpson indices were calculated for all three methods but were largely uncorrelated including between conventional methods. Although the technology is relatively new, optical sensors may provide next-generation insight into the spatiotemporal dynamics of invertebrate biodiversity and their conservation.

**Significance Statement:** The implications of this research extend from the field level to the regional level. Much of what scientists understand about the decline of invertebrates comes from a small number of long-term studies that can be coarse and correlational in nature. High-resolution biodiversity data sets on fields to landscapes may provide the insight needed for the successful management and accounting of biodiversity by practitioners and policymakers. Such high-resolution data has the potential to support global efforts and coordination of biodiversity conservation.

## Introduction

Invertebrate species biodiversity is fundamental to ecosystem processes, functions, and services (Yang & Gratton, 2014). Monitoring metrics of invertebrate biodiversity including species richness, abundance, evenness, and biodiversity indices (e.g. Shannon index) can inform management and policy at multiple scales. Such data are critical to agricultural production and sustainability (Landis, 2017). However, invertebrate biodiversity metrics are difficult to quantify (Geiger et al., 2016; Shortall et al., 2009) and monitor at broad spatial and temporal scales (Sánchez-Bayo & Wyckhuys, 2019; Tilman et al., 1994). The difficulty is largely due to the necessity of skilled labor required for taxa identification on which biodiversity quantification relies (Wägele et al., 2022), and is both limited and prohibitively costly (Gardner et al., 2008). Common approaches to collecting insect inventories include sweep netting as well as Malaise-, pan-, and light traps. Each method has its own bias toward certain insect groups (Bick et al., 2020) often resulting in the concurrent use of techniques in studies and practice (LaCanne & Lundgren, 2018).

New technology is needed to monitor invertebrate biodiversity in real time for agricultural systems. Such a tool would provide data to support biodiversity-focused management at field to landscape scales (LaCanne & Lundgren, 2018) and allow for tracking of the impact of conservation measures, or the lack thereof. Automation of systems has the potential to reduce labor, time, costs, and human error. While many automated insect monitoring tools are available for agricultural pest monitoring (Bick et al., 2023; Silva et al., 2013), overall, these approaches are not suitable for assessing biodiversity as they focus on the identification of indicator species, not communities (J. G. Lundgren & Fausti, 2015a). The automatic quantification of invertebrate biodiversity could improve the data available for monitoring and evaluation of conservation efforts but currently, no method exists at scale (Wägele et al., 2022), despite calls for such data and analytics to inform the assessment and management of ecosystems (García et al., 2023).

Real-time data on invertebrate biodiversity likely would improve our understanding of insect population changes at a regional or even global scale, filling a gap in the tracking of insect change. The incorporation of ‘big data’ has been shown to help mitigate some methodological biases (Geiger et al., 2016). One such effort is the global malaise project that is using automated taxonomic identification from traps using DNA, addressing the most labor-intensive part of this method (Krishna Krishnamurthy & Francis, 2012). It is a highly promising ‘big data’ approach; unfortunately, the method over-represents known species, has an inherent sampling bias towards flying insects and emphasizes species with large mitochondrial differences. Optical entomological methods such as lidar, where an optical signal is recorded from insects flying through a beam of emitted light, can record large numbers of insect flights without using a lure. However, it is unclear how optical sensors compare to conventional methods in measuring populations and communities (Garcia et al., 2023; Rydhmer et al., 2022).

The goal of this study is to determine if the measurement of an insect biodiversity metric can be automated with the use of optical near-infrared insect sensors. In this work, we deployed sensors (Rydhmer et al., 2021) in 20 agricultural fields across six crops in Kansas, USA. The sensors were deployed alongside Malaise traps and the sites were sampled with sweep nets. Each site was evaluated on two different occasions to capture seasonal changes. Specifically, we compared conventional methods to each other and with the automated biodiversity metric utilizing unsupervised clustering of data collected by a lidar-based sensing method.

## Materials and Methods

### Data collection

Insect populations were monitored at 20 sites (Figure 1) in June and July of 2020 using sensors alongside conventional methods (Malaise traps and sweep nets) to evaluate sensor-based versus conventional metrics of biodiversity. Representative agricultural crops of central Kansas were sampled including three corn, three sorghum, six soybean, one alfalfa, two pasture, and five complex cover crops. The complex cover crops consisted of approximately eight species of annual grass and forb cover crops.

**Figure 1.**
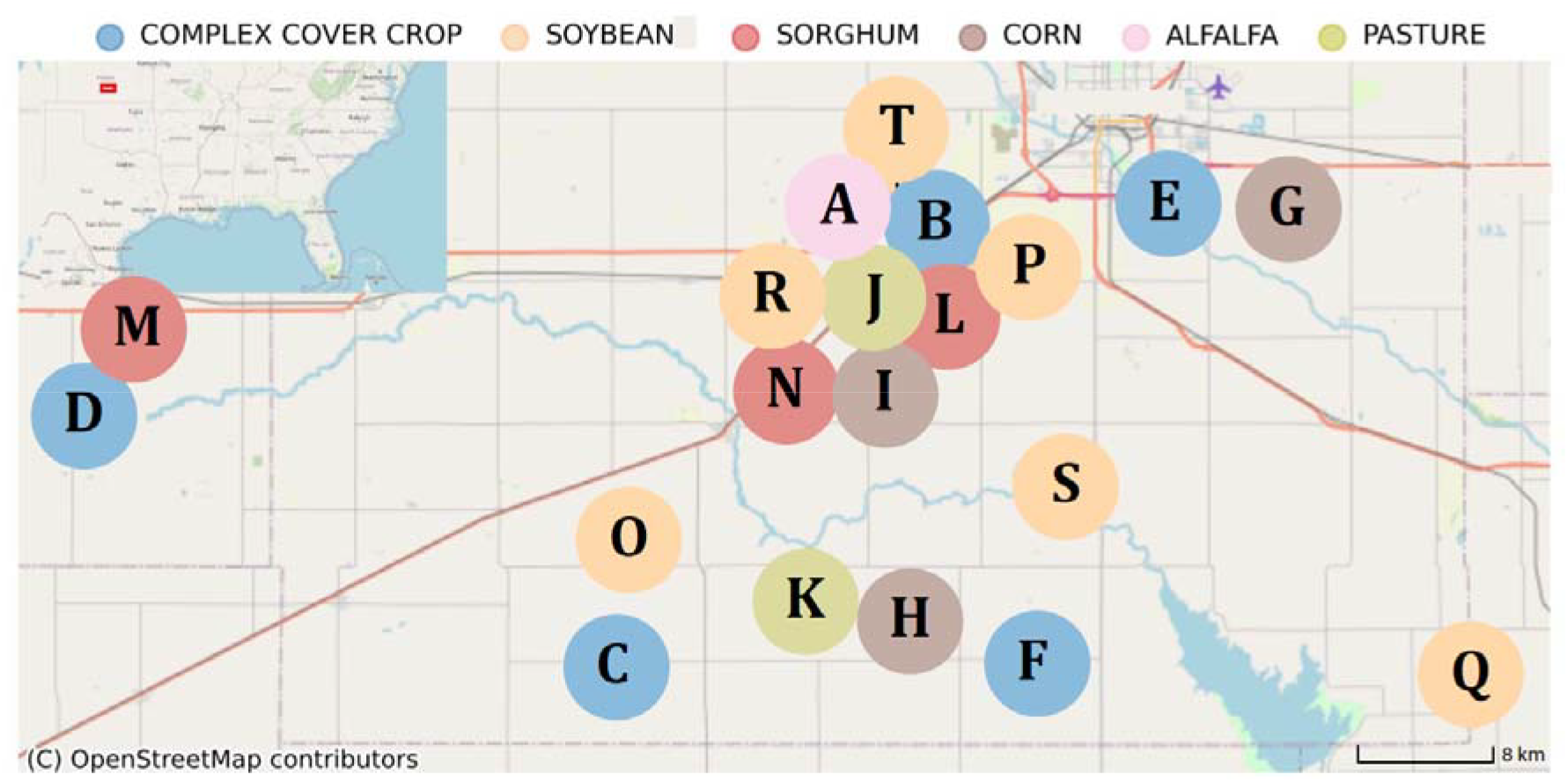
Map of 20 field site locations distributed around central Kansas. Fields are color-coded by crop type. Field dots are enlarged and shifted to maintain the anonymity of the participating farms. Map data from www.openstreetmap.org.

**Figure 2.**
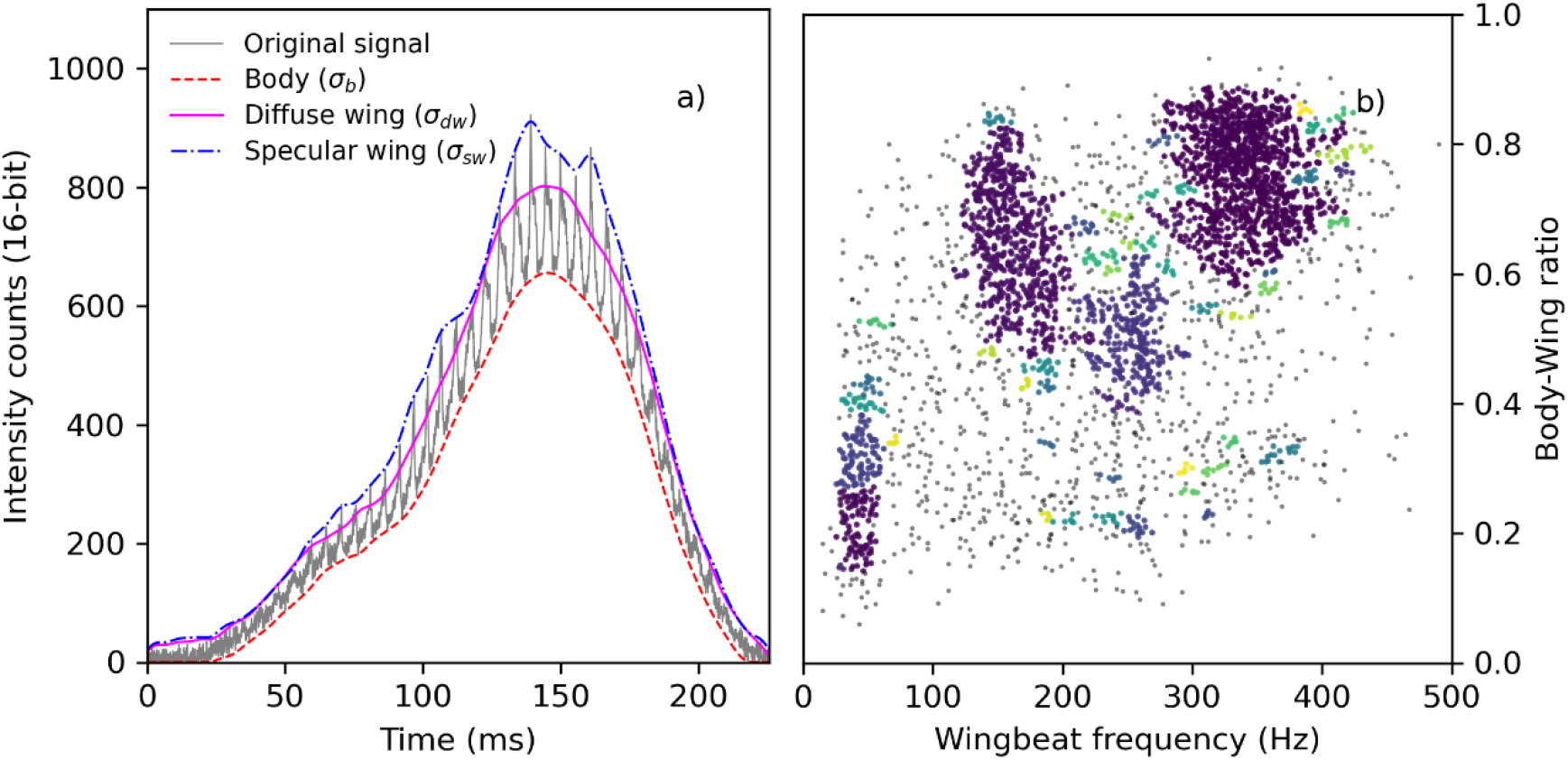
Example of an insect event’s signal and clustering. a) An example of an insect event recorded from the sensor. The wing beats are visible as modulations on top of the signal. The dashed red, solid magenta and dash-dotted blue curves show the body, diffuse- and specular wing signals respectively. The BWR is the ratio between the magnitude of the body- and diffuse wing signal. b) Clustered insect events recorded by the sensor in a soybean field (Field R) in July. The grey events are too sparse to form clusters and are therefore discarded.

An autonomous near-infrared sensor (described in (Rydhmer et al., 2021) and produced by FaunaPhotonics ApS., Copenhagen SV, Denmark) was placed ∼50 m from the field margin and was monitored continuously for two periods of three days in June and in July. The sensor uses light-emitting diodes to transmit infrared light (810 nm & 970 nm), creating a measurement volume between 5 and 70 L, depending on insect size (Rydhmer et al., 2021). Insects flying in front of the sensor back-scatter light, which is recorded by a photodiode as a time signal. Insect recordings are automatically separated from noise originating from other sources (e.g. rain drops or plant interference) using proprietary cloud-based neural network software, as used in Bick et al., 2023 and Rydhmer et al., 2021. Additionally, observations without clearly identified wingbeats or body-to-wing ratios were discarded. A total of 1,057,115 observations were recorded, of which 106,083 remained after filtering and were included in the study. A recorded observation consists of time series data from which information pertaining to the physical features of the individual insect can be obtained (Rydhmer et al., 2021). Sensors were compared with conventional sampling of invertebrates (sweep nets and Malaise traps) in the same fields. Foliar and low flying insects were captured using a sweep net (38 cm diam., Bioquip™, Rancho Dominguez, CA, USA). Insects were collected at 50, 100, and 150 m from the field edge along a linear transect. Sweeps (n = 50 per location) were performed perpendicular to the transect, parallel to the field edge. Insects were transferred to a sealed plastic bag and were frozen until sorted from the plant material and identified in the laboratory.

Malaise traps were deployed at each site to capture the aerial insect community. A single bi-directional, Townes-style trap (dimensions 1.8 long; 1.8 m at its tallest height, and 1.2 m at its shortest height) was placed 100 m from the margin and adjacent to the ecosystem service sampling areas. The wall of the trap was parallel with the field margin. The traps were allowed to operate for 24 h, and the insects captured in the collection vials were preserved in ethanol.

All specimens collected by sweep net and malaise traps were identified to the lowest possible taxonomic unit (i.e., species or morphospecies). Due to a lack of species identification knowledge and time limitations, thrips (Insecta: Thysanoptera) were not identified beyond the family level and were not included in community metrics analyses (abundance, species richness, and diversity). All immature insects were identified to family and grouped together, except for lepidopteran larvae, which were categorized as morphospecies independent of the adult stage due to their functional differences. All other specimens were identified to species using written and online taxonomic keys. Specimens that were not able to be positively identified to species were separated into distinct morphospecies. Voucher specimens of all taxa are housed in the Mark F. Longfellow Biological Collection at Blue Dasher Farm, Estelline, SD.

Invertebrate predator abundance and predation rates of insect and weed seeds were monitored to evaluate ecosystem services in each field. The abundance of invertebrate predators was determined by identifying and quantifying all predators from the soil and foliar samples. Predation rates in each field were assessed by establishing 15 sentinels wax moth larvae (*Galleria mellonella* L. [Lepidoptera: Pyralidae]) per plot arranged in three 5 × 3 7.5 m grid orientations (n = 45 per field) following Lundgren et al., 2006 protocol. Weed seed predation by invertebrates was assessed for soil and foliar granivore communities using seed cards as described in Lundgren et al., 2006. Granivory were measured on three abundant weed species (Johnsongrass (*Sorghum halapense* (L.) Pers.; Poaceae), lambsquarters (*Chenopodium album* L.; Amaranthaceae) and redroot pigweed (*Amaranthus retroflexus* L.; Amaranthaceae), V & J Seed Farms, Woodstock, IL, USA). Seeds were attached to 10 × 8 cm plastic cards (Avery™ insertable plastic dividers; #11200; Brea, CA, USA) using 6 cm strips of double-sided tape (Scotch, 3M, St Paul, MN, USA). Each species (n = 20 seeds of each species; 60 seeds per card) were placed on a 2 × 10 pattern on each card. Fine quartz sand was spread over exposed areas of the tape to exclude visiting invertebrates. To exclude granivorous vertebrates, a wire cage (14 × 12 cm cage, 1.4 × 1.4 cm mesh opening) was placed over the card and placed >3 cm above it. Control cards were used to account for seed loss from environmental factors such as wind or rain and contained 1.5 × 1.5 mm black glass beads (Cousin™ DIY, #AJM61215021, Largo, FL, USA) of comparable size as the weed seeds (Lundgren et al., 2006). Each plot received three seed cards and one control card (n = 9 seed cards and three control cards per field), placed on the soil surface in the four corners of each plot. Granivory was measured as the number of seeds removed or damaged per card after a 3-day exposure.

### Data analysis

The wing-beat frequency (WBF) and body-to-wing ratio (BWR) were calculated from all observed insect flights similarly to previous groups (Gebru et al., 2018; A. Genoud et al., 2020; A. P. Genoud et al., 2019). The signal from the insect body (σ_b_) and the diffuse and specular signal contributions from the insect wings (σ_dw_ and σ_sw_) are estimated and separated using sliding minimum, sliding median and sliding maximum filters with a filter width corresponding to the wing beat period of the insect. The BWR is defined as the closed ratio between the body and wing contributions according to equation (1). An example of an insect signal is shown in Figure 1a.

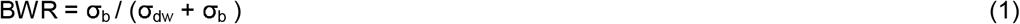

Insects of the same species exhibit similar physical properties, and therefore also similar signal features (Kirkeby et al., 2021). Normalization of the feature space is a standard procedure prior to clustering. While BWR values are bound between 0 and 1 by definition (equation 1), WBF values frequencies typically vary between 20 Hz and 1 kHz (Jansson et al., 2019). WBFs were therefore divided by 1000 to produce values falling predominantly between 0 and 1. For clustering, we used the DBSCAN (Density-based spatial clustering of applications with noise) algorithm (Ram et al., 2010) due to its suitability in identifying clusters without a Gaussian distribution assumption (Ester et al., 1996). DBSCAN uses two parameters, the minimum number of insects needed to form a cluster (min_samples) and the merge distance □, to determine which observations to merge into clusters. Data points too far away from any cluster and too sparsely distributed to form a new cluster are defined as outliers. This method was used to calculate the number of clusters or distinct groups (i.e. richness) and a diversity index of cluster groups based on Shannon and Simpson indices.

All insects collected with Malaise traps and sweep nets were classified by order, family, and species when possible. Then species richness (defined as the number of distinct taxonomic species present, independent of abundance), Shannon index, and Simpson index were calculated on the insect samples from both conventional methods for each field in June and July.

The data from the capture methods were randomly divided into two data sets: one used to optimize the DBSCAN clustering algorithm, and the other used for testing. To have a sufficiently large test set, the optimization set was limited to 30% of the data collected. During the optimization, _LJ_ and min samples were tuned to maximize the Spearman correlation between biodiversity metrics from the sensors and conventional metrics using stochastic gradual descent. This process was repeated for the richness and Shannon and Simpson indices for each of the trapping methods, plus an additional model fitted to the combined species richness from both conventional methods. Shannon and Simpson indices were not calculated on the combined dataset since these indices rely on the relative abundance of species, which are not comparable between the two methods.

Optimal parameters could be found that produced significant correlation (p < 0.05) for four of the seven comparative measures; however, no parameters could be found that satisfactorily modeled the Shannon index from the sweep netting nor the Simpson index for either trapping method.

Spearman-rank correlations between the clustering results calculated from the optical sensor data and the biodiversity measures obtained with the two physical insect field-sampling methods were calculated. Additionally, Analyses of Variance (ANOVA)and TukeyHSD post hoc analyses were conducted to evaluate the impact of sampling month, crop type, and field on richness estimates.

## Results

In total, 106,083 insect flight events were recorded by the sensors. The Malaise traps collected 14,641 insects, whereas sweep nets collected 15,858 insects (Figure 3). The optical sensors recorded approximately one order of magnitude more insect flights compared to the number of insects collected with each of the conventional methods (Figure 3). Insect abundance was uncorrelated between all methods, including both conventional methods and the automated method (Figure 4; Malaise trap counts and sweep net counts r = 0.25, p =0.16, sensor events and sweep net counts r = 0.05, p = 0.78, sensor events and Malaise trap counts r = 0.05, = <0.88).

**Figure 3.**
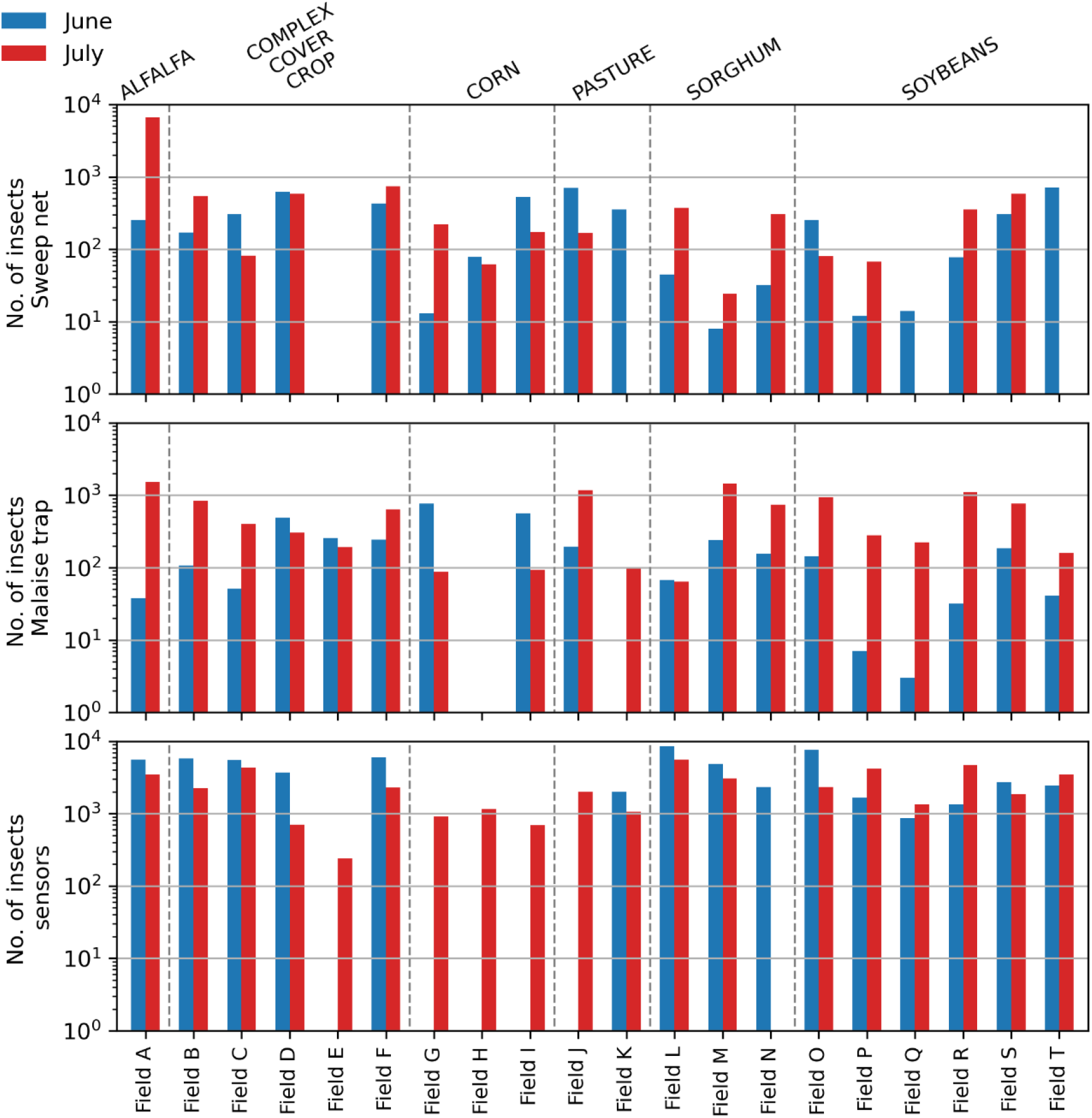
The number of insects collected using sweep nets (top panel) and Malaise traps (middle panel), and insect flight events recorded with the sensor (bottom panel) per field. Insect numbers are separated by month with insects observed in June visualized with blue bars and in July with red bars.

**Figure 4.**
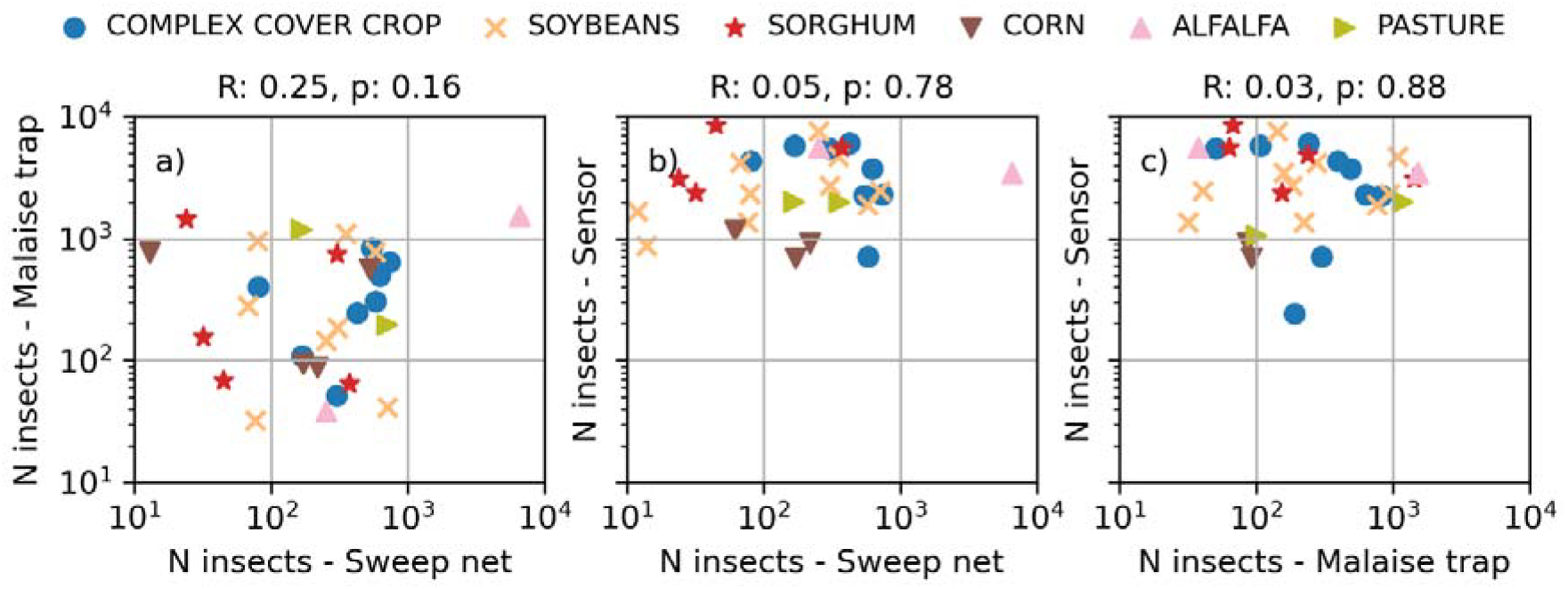
Scatter plots of measured insect abundance comparing the monitoring methods on a logarithmic scale. a) Scatter plot of the number of insects captured with sweep nets and Malaise traps. b) Scatter plot of the number of insects captured with sweep nets and the number of insect flight events recorded by the sensor. c) Scatter plot of the number of insects captured with Malaise traps and the number of insect flight events recorded by the sensor. No correlations were found on insect abundance for any of these methods.

Comparing the relative insect abundance between orders collected with conventional methods (Figure 5) depicts differences in capturing biases due to methodology. Diptera were most frequently collected from the Malaise traps, whereas Hemiptera, then Diptera and Psocoptera were more frequently captured with sweep nets. In general, less flight-active insects were more prominent in the sweep net data.

**Figure 5.**
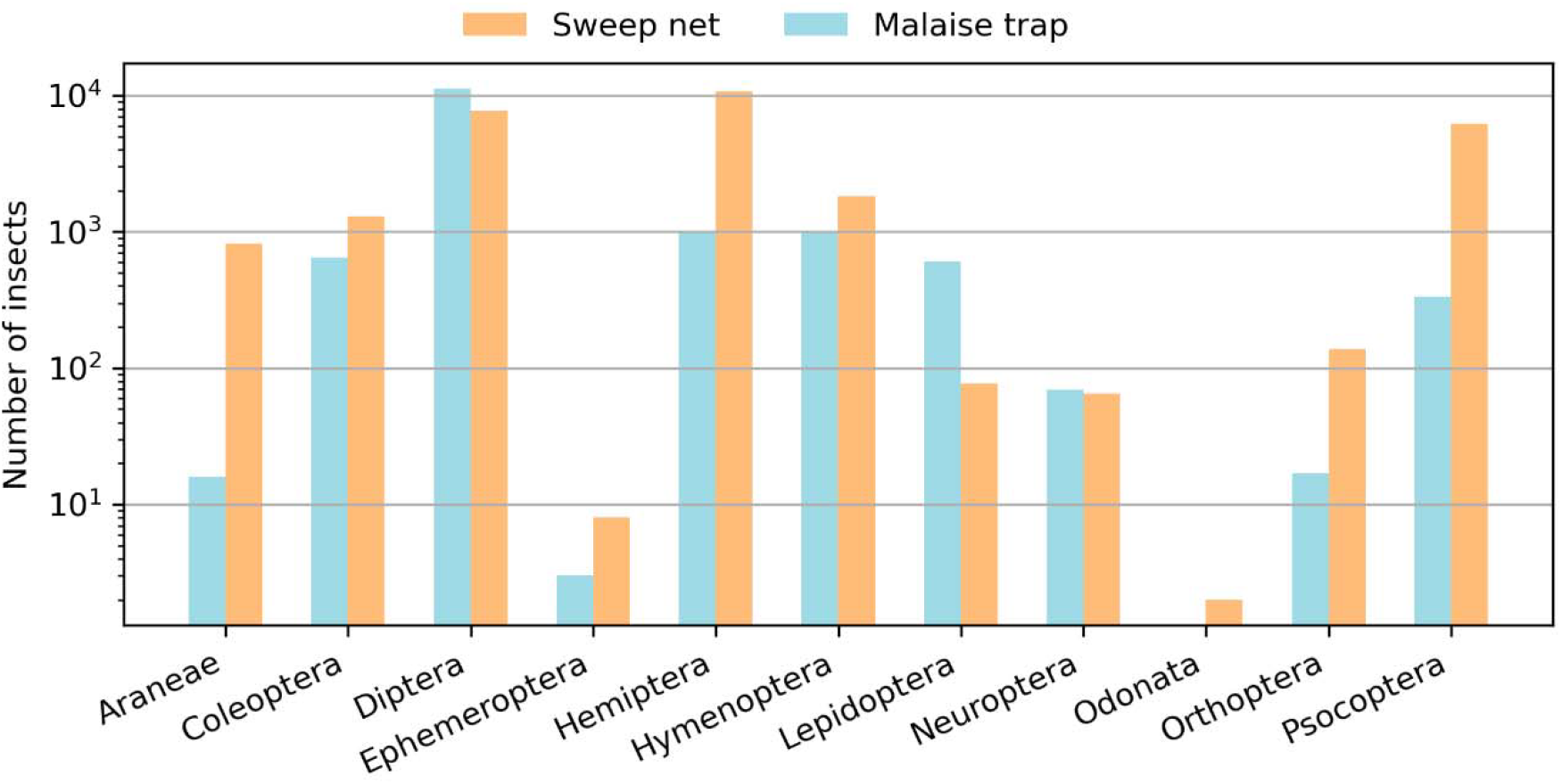
The number of insects collected with sweep net sampling and Malaise trap monitoring, aggregated by order.

There were no discernible differences in variation between time points from sensors (F = 6.091, Pr = 0.0191) and sweep nets (F = 1.326, Pr = 0.258). However, Malaise trap abundance showed significantly greater insect densities in July (*µ* = 76.9, F = 9.71, Pr = 0.0037) than June (*µ* = 43.3). Crop type was found to impact sweep net abundance (F = 3.369, Pr = 0.008) but not sensor (F = 1.644, Pr = 0.17) or Malaise trap (F = 1.692, Pr = 0.152) abundance estimates. A series of TukeyHSD post hoc analyses found no differences in abundance estimates between sample time points for each field.

Insect species richness was estimated from sensor-recorded insect flight events using a set of seven DBSCAN parameters (models) to cluster the held-out test data, yielding the number of clusters per field – the novel richness metric (Figure 2b). The correlation between the number of clusters and each of the comparative biodiversity metrics are shown in Table 1.

**Table 1.** Correlations between the automated biodiversity metrics calculated from sensed insect data, and those obtained from Malaise trap and sweep net collections. Rows in the table denote which data was used to fit the clustering algorithm, whereas columns indicate which parameters the obtained correlations refer to. Correlations with a p-value below 0.05 are significant and marked in bold.

| Fitting data | Correlation coefficients (R, p-value) |  |  |  |  |  |  |
| --- | --- | --- | --- | --- | --- | --- | --- |
|  | Richness |  |  | Shannon |  | Simpson |  |
|  | Malaise trap | Sweep net | Combined | Malaise trap | Sweep net | Malaise trap | Sweep net |
| Richness Malaise | **0.42,**<br>**0.050** | 0.26,<br>0.246 | 0.37,<br>0.112 | **0.44,**<br>**0.044** | -0.22,<br>0.359 | 0.35,<br>0.114 | 0.01,<br>0.964 |
| Richness sweep net | **0.45,**<br>**0.035** | **0.44,**<br>**0.046** | **0.47,**<br>**0.035** | 0.11,<br>0.612 | 0.20,<br>0.378 | 0.09,<br>0.702 | 0.16,<br>0.475 |
| Richness combined | **0.52,**<br>**0.014** | **0.48,**<br>**0.028** | **0.55,**<br>**0.012** | 0.21,<br>0.344 | 0.20,<br>0.394 | 0.14,<br>0.534 | 0.14,<br>0.537 |
| Shannon Malaise | **0.42,**<br>**0.050** | 0.26,<br>0.246 | 0.37,<br>0.112 | **0.44,**<br>**0.044** | -0.22,<br>0.359 | 0.35,<br>0.114 | 0.01,<br>0.964 |
| Shannon sweep net | **0.45,**<br>**0.037** | 0.37,<br>0.097 | **0.46,**<br>**0.040** | 0.27,<br>0.221 | 0.22,<br>0.339 | 0.17,<br>0.459 | 0.08,<br>0.733 |
| Simpson Malaise | **0.44,**<br>**0.041** | 0.32,<br>0.159 | 0.43,<br>0.061 | -0.16,<br>0.484 | 0.03,<br>0.893 | 0.02,<br>0.913 | 0.06,<br>0.797 |
| Simpson sweep net | 0.05,<br>0.837 | -0.13,<br>0.569 | -0.07,<br>0.774 | **-0.73,**<br>**0.005** | 0.07,<br>0.817 | -0.04,<br>0.846 | 0.12,<br>0.598 |

Per field, the mean number of clusters that approximate richness was 41.1 (N = 34, SE = 3.29). The Malaise traps had a mean field richness of 60.5 species (N = 37, SE = 6.43) containing 10 orders, 146 families, and 709 species. The mean richness observed in the sweep nets was 47.4 species (N = 35, SE = 5.53), containing 11 orders, 149 families, and 664 species. Combined, the collected samples with both field-sampling methods contained 941 species distributed over 183 insect families in 11 orders.

The three models fit on species richness are generally comparable (fit on sweep net, Malaise trap, and their combined data). Identical DBSCAN parameters were calculated when the models were fit on the combined sweep net and Malaise trap richness. We therefore used this model termed the ‘automated biodiversity metric’ to evaluate the relationships between sensors, and the conventionally measured species richness and ecosystem services.

All species richness metrics were correlated (Figure 6a-d). The weakest correlation was between Malaise trap and sweep net richness metrics (R = 0.36, p = 0.046). The correlation between the number of clusters calculated in the sensor data and the conventional models was strongest for the combined richness, which was what the model was fitted on (R= 0.55, p = 0.012; Figure 6d). Significant yet weaker correlations were also found between the model and the Malaise trap and sweep net richness respectively (R = 0.52, p = 0.014; R = 0.48, p = 0.028).

**Figure 6.**
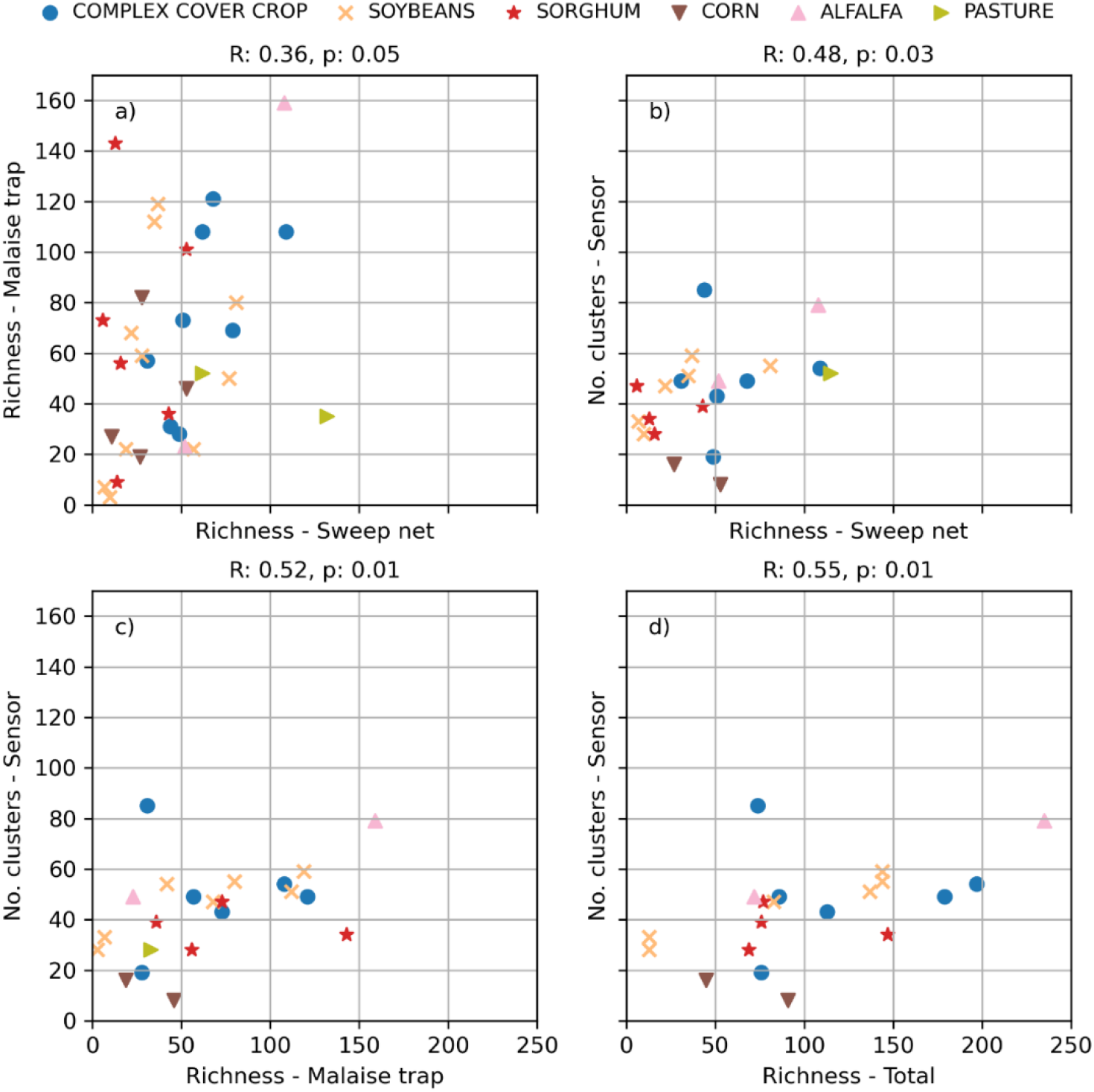
Scatter plots and Spearman correlations for the species richness estimations across all models. The sensor results are from the model fitted to the total richness in both Malaise traps and sweep nets. a) Species richness calculated from Malaise traps vs. sweep net samples, b) species richness calculated from sensor clusters vs. sweep net samples, c) species richness calculated from sensor clusters vs. Malaise trap samples, and d) species richness calculated from sensor clusters vs. total richness across traps and sweeps.

No correlations were found when comparing sensor richness to any ecosystem services (Table 2). Conventional sampling methods were typically not correlated with ecosystem services with one exception. Sweep net species richness was positively correlated with the percent of waxworms predated (Table 2).

**Table 2.** Spearman’s correlation table between richness metrics calculated from sweep nets, Malaise traps, combined conventional methods, and sensor clusters (automated biodiversity metric) compared to ecosystem services of percent waxworm predation, total number of predators, Johnsongrass predation, Pigweed predation, Lambsquarter predation, and all seed predation.

| Richness metrics | % Waxworms Predated | Total # of predators | Johnsongrass predation | Pigweed predation | Lambsquarter predation | All seed predation |
| --- | --- | --- | --- | --- | --- | --- |
| Spearman's correlation coefficient (R, p-value) |  |  |  |  |  |  |
| sweep net | **0.49, 0.04** | 0.34, 0.17 | -0.37, 0.13 | 0.05, 0.85 | -0.04, 0.89 | -0.07, 0.78 |
| Malaise trap | -0.16, 0.52 | -0.13, 0.61 | -0.16, 0.52 | -0.36, 0.15 | -0.40, 0.10 | -0.45, 0.06 |
| total - richness | 0.21, 0.43 | 0.16, 0.56 | -0.39, 0.14 | -0.26, 0.33 | -0.32, 0.23 | -0.41, 0.11 |
| sensor clusters | -0.11, 0.66 | -0.26, 0.29 | 0.14, 0.56 | -0.11, 0.66 | -0.15, 0.54 | -0.11, 0.66 |

## Discussion

This work serves as the first field-validated insect biodiversity metric using autonomous distributed optical sensors (Kouakou et al., 2020). The automated biodiversity metric was calculated and validated from flight events of sufficiently high quality to be able to extract a wingbeat frequency and body-wing ratio. Our results suggest that the sensor-derived metric is correlated with conventional estimates of biodiversity. Specifically, the sensor-derived biodiversity metric optimized for insect species richness is more correlated with each conventional sampling method than these methods are to one other (Figure a-d) (Malaise trap – sweep net R = 0.36; Malaise trap – sensor R = 0.52; sweep net – sensor R = 0.48). This indicates that metrics derived from optical sensors are likely generalizable and thus have the potential to provide accurate and autonomous measurements of insect species richness. Ecosystem generalizability was demonstrated by deriving and testing the biodiversity metric across six major crops in central Kansas. Still, future work is needed to evaluate the extent to which the metric may be generalized across agroecosystems outside our study area and to other terrestrial ecosystems. Current results indicate that the insect diversity metric may be calculated for a variety of functionally different crops without the need to classify insects into taxonomic groups. This approach is beneficial as such classification at present requires skilled labor and significant time. This metric may provide new insight into the management of ecosystems as significant and growing evidence suggests that biodiversity is correlated with greater ecosystem functions such as pest control (Lundgren & Fausti, 2015). Future work may focus on characterizing the composition of insect communities and species to address specific needs of managing ecosystems beyond biodiversity.

The automated biodiversity metric and Malaise trap species richness significantly correlated across all method iterations, save one (Table 1). Sweep net richness only correlated with two of the method iterations. The stronger correlation between the sensors and Malaise traps is hypothesized to result from these methods monitoring flying insects continuously, compared to sweep net sampling. Results were less clear for the correlations with Shannon and Simpson species diversity indices. The models fitted on Malaise trap richness also was significantly correlated with for Malaise trap Shannon index (Table 1). This is likely due to the co-correlation between the richness and Shannon index in the malaise trap (R=0.6, p=0.01, Supplementary Table 3). Other similar curiosities, such as the negative correlation with the Malaise trap Shannon index achieved when fitting on the sweep net Simpson index are also assumed to be the results of co-correlations between the conventional methods. A full table of all co-correlations is included as supplementary material (Supplementary Table 3). However, when fitting on Shannon index from the sweep-net data no correlation was found. This is likely due to overfitting, where the model performed well on the fitting data but did not generalize to the rest of the dataset. A larger fitting dataset is needed to resolve this issue. No model resulted in significant correlations in Simpson indices between any of the sampling methods. The lack of consistent correlations between biodiversity metrics may also reflect the nature of the biodiversity indexes which considers species evenness, a characteristic not fully accounted for in the DBSCAN algorithm based on minimum thresholds for classification of clusters.

The sensor observed the greatest number of insects, recording almost one order of magnitude more than both the Malaise traps and sweep nets (7.25 and 6.69 times, respectively). This difference is likely explained first by the observational period and then by methodology. Both sensors and Malaise traps continuously monitored each field, unlike sweep nets which collect insects at discrete time points. The sensors’ monitoring period was three times longer than the Malaise traps. However, even after accounting for the greater measurement period, the sensor methodology was still 2.42 times more efficient than Malaise traps at observing all insects. A previous study reported sensors observed 19 times the number of insects compared to those collected in water traps, another continuous monitoring method (Rydhmer et al., 2021). Specific insect species are also detected more efficiently. For example, the sensor was reported to be 18.6 and 6.7 times more efficient than plant counts (discrete sampling) and water traps (continuous monitoring) at observing pollen beetles (*Brassicogethes aeneus*; Bick et al., 2023). Greater sampling efficiency may be associated with measurement volume, a potential correlate of insect counts. Optical sensors with greater measurement volumes report recording tens of thousands of insect flights per day (Brydegaard et al., 2020). A potential confounding effect of sensors is the possible ‘double counting’ bias, as an individual insect can be recorded repeatedly. While ‘double counting’ possibly explains the greater number of observed insects, such biases are a common limitation of count-based inferences on population dynamics (Elphick, 2008). The Law of Large numbers indicates that the greater insect observations will likely be more representative of the ecosystem (Balm & Katz, 1965). Despite its potential limitations, our understanding of complex species and community dynamics can benefit greatly from automation that significantly increases sampling intensity across space and time.

The lack of correlation of abundance across all three methods (Figure 4) is surprising as previous work has shown correlations between sensor measurements and water traps for insect abundance (Rydhmer et al., 2021). Disparities in sampling timing may be contributing to the lack of correlation. While sweep netting occurred in conjunction with setting up or taking down the Malaise traps, these efforts were substantially less correlated with the setup of the optical sensors: to the nearest 3 days in June and the nearest 22 days in July. The lack of correlation between the Malaise traps and the sensors may be due to the long period between the monitoring sessions at each site. Insect flight activity is heavily influenced by the weather, or the seasonal differences between the beginning and end of July – both of which may also explain the significance of the month on Malaise trap data. An additional factor may be the high noise composition of the recorded signals due to plant interference. During cleaning of this dataset, it is possible that variations in the relative degree of noise signals between fields (e.g. as a result of different crop heights and stiffness) resulted in more data loss from noisier fields, thus introducing a systematic error in abundance measurements for the sensor data. However, it should be noted that we observed no statistical correlation in abundance collected with Malaise traps versus sweep nets.

One challenge with the sensor’s dataset was the high proportion of noise signals, thought to result from plant interference. Of the total 1,057,115 signals recorded by the sensor, only ∼10% were classified as insects and included in the analysis. While we believe the noise classification filter is highly accurate, misclassified events may alter the total count. The signals generated by insects and plants moving through the sensor’s measurement volume are very different. Most non-insect events show no high-frequency components and are therefore correctly removed by the noise filter. However, plants modulating in front of the sensor may appear to have a wing beat frequency and would be misclassified as insects. It is also hypothesized that strong signals generated by vegetation interference may obscure weaker signals generated by small insects. Regardless, misclassification is likely low enough to substantially alter the representation of the insect population or the automated biodiversity metric.

Autonomous optical sensors, such as the ones used in this study, provide continuous, potentially real-time monitoring to support next generation, ‘big data’ insight to the field of entomology (Rosenheim and Gratton 2017). Currently available ‘big data’ sources provide spatio-temporally rich environmental and management data but may require similarly rich spatio-temporal data sets on the population and community dynamics of invertebrates to fully enable new insights promised by ‘big data’ technology. Autonomous sensors for monitoring biodiversity provides such data to support ‘big data’ analytics. The use of sensors for insect species richness monitoring is faster, in this case potentially more representative of richness, and likely cost-effective due to a decrease in labor compared to conventional methods. Sensors complement conventional sampling methods by allowing for real-time estimation of biodiversity, reducing time lags associated with traditional species inventory. Automated methods presented in this paper, once a generalizable calibration has been determined, offer faster estimates of biodiversity which will support time-critical decision making and conservation planning efforts. Methods such as the one described in this paper do not rely on identifying taxonomic groups and remove human error (a major concern for insect identification). Furthermore, standardized sensors are not prone to local and regional variations in sampling methods and may therefore be able to facilitate comparative biodiversity monitoring on a global scale.

There is a need to scale up and scale down sensor monitoring to understand species dynamics. The current technology complements the entomological radar group ‘BioDAR’ which is aiming to use libraries of insect radar signals for functional group classifications for high flying migratory insects at a regional scale (Rhodes et al., 2022). Similar estimates of insect functional groups might be similarly inferred from optical sensor recordings for all flying insects on a field scale.

Similarly, vertical looking radar is used to classify insects into higher level taxonomic groups such as Order or even Genus (Chapman et al., 2002; Stefanescu et al., 2013; Wood et al., 2009). It seems likely that similar or even higher precision can be achieved by including taxonomic information in clustering algorithms, such as specific orders (e.g. Lepidoptera, Coleoptera, Diptera). Future work could focus on identifying these groups, determining functional biodiversity, and quantifying their contribution to ecosystem services.

The current study shows a single instance of correlation between richness and a measure of ecosystem services. Greater species richness does not always translate into an increase in functional biodiversity or ecosystem services, as there is often ecological redundancy (Greenop et al., 2018). The lack of a relationship may also reflect different ecological interactions among species in the upper canopy versus above canopy level. These questions can be further explored in future work when the sensor’s ability to estimate functional biodiversity has been developed.

## Conclusion

Conservation of biodiversity is gaining recognition as a global challenge with similar significance to climate change. However, unlike global climate, species populations and biodiversity function across different local to regional spatiotemporal scales. Detailed data on insect diversity across these scales is needed to assess the decline and inform conservation efforts. There is a call to automate the collection of *in situ* data and integrate such data with remote sensing-based models to accelerate conservation of global ecosystems (Garcia et al 2023). These integrated technology and big data approaches are especially needed for the conservation of invertebrates and could expand upon and accelerate the long-term and detailed monitoring efforts of invertebrate biodiversity (Sánchez-Bayo & Wyckhuys, 2019) The current study demonstrates successful development of *in situ* data collection able to be integrated with remote sensing models as described by Garcia et al 2023. Such approaches are poised to support the Intergovernmental Science-Policy Platform on Biodiversity and Ecosystem Services with rich, real-time data and help inform global biodiversity models that have had to rely on coarse, low-resolution data sets in some cases (Schipper et al., 2020).

Real-time feedback of simple metrics of biodiversity would greatly benefit agriculture by demonstrating the association between management and biodiversity in real-time, without time lag to process samples. The combination of pest detection with optical sensors (Bick et al., 2023; Kirkeby et al., 2021) and monitoring of biodiversity may inform integrated pest management and reduction of pesticide applications. Farming informed by such data analytics may deliver significant benefits to farmers including substantial reductions in pests (Lundren and Fausti 2015) and significant economic benefits at both farm and regional levels (Landis et al 2008, LaCanne and Lundgren 2018). Networks of local monitoring sensors may scale up to infer regional biodiversity and inform its management. The current technology could complement the global Malaise trap initiative by increasing the number of observations by an order of magnitude and providing earlier warning signs of regional pest movement or species decline. Our results suggest AI supported human expertise may provide the most efficient, robust inference on biodiversity.

Thus, we are not advocating such technology replaces conventional monitoring, but rather that this automation enhances the state of the art. Autonomous monitoring has the potential to revolutionize the field of entomology by forming the basis for a next generation of species to community insect models predicting the dynamics of invertebrates.

## Acknowledgments

We thank the USDA Cheney Lake Conservation District (Lisa French), Understanding Ag (Ray Archuleta) for logistical support and field assistance. Tom Rabaey (General Mills) provided helpful feedback and advice to guide this project. We thank Kevin James Knagg and Mads Fogtmann from FaunaPhotonics A/S for facilitating this work. We thank Ecdysis Foundation field and lab technicians for collecting, processing, and Dr. Kelton D. Welch for identifying invertebrate specimens. We thank the farmers cooperating in the General Mills regenerative Agriculture Programs who granted us access and permission to sample their farm fields.

## Supplementary information

**Supplementary Table 1.** A table describing the crop type and number of insects observed in each field in June and July across all three methods.

| Field name | Crop type | Sensor |  | Malaise trap |  | Sweep net |  |
| --- | --- | --- | --- | --- | --- | --- | --- |
|  |  | June | July | June | July | June | July |
| A | ALFALFA | 5529 | 3437 | 38 | 1530 | 254 | 6579 |
| B | COMPLEX COVER CROP | 5739 | 2244 | 108 | - | 79 | 62 |
| C | COMPLEX COVER CROP | 5454 | 4290 | 51 | 828 | 169 | 546 |
| D | COMPLEX COVER CROP | 3718 | 704 | 493 | 93 | 525 | 172 |
| E | COMPLEX COVER CROP | - | 240 | 258 | 87 | 13 | 219 |
| F | COMPLEX COVER CROP | 6008 | 2281 | 243 | 303 | 626 | 583 |
| G | CORN | - | 912 | 761 | 278 | 12 | 68 |
| H | CORN | - | 1158 | - | 398 | 303 | 81 |
| I | CORN | - | 687 | 561 | 636 | 427 | 740 |
| J | PASTURE | - | 1993 | 195 | 1437 | 8 | 24 |
| K | PASTURE | 1980 | 1055 | - | 64 | 45 | 375 |
| L | SORGHUM | 8492 | 5545 | 68 | 743 | 32 | 304 |
| M | SORGHUM | 4845 | 3080 | 239 | 948 | 254 | 80 |
| N | SORGHUM | 2349 | - | 155 | 1176 | 708 | 167 |
| O | SOYBEAN | 7524 | 2310 | 144 | 101 | 358 | - |
| P | SOYBEAN | 1657 | 4162 | 7 | 225 | 14 | - |
| Q | SOYBEAN | 865 | 1349 | 3 | 1090 | 77 | 354 |
| R | SOYBEAN | 1349 | 4679 | 32 | 160 | 713 | - |
| S | SOYBEAN | 2696 | 1867 | 184 | 771 | 307 | 580 |
| T | SOYBEAN | 2443 | 3442 | 41 | 192 | - | - |

**Supplementary Table 2.**
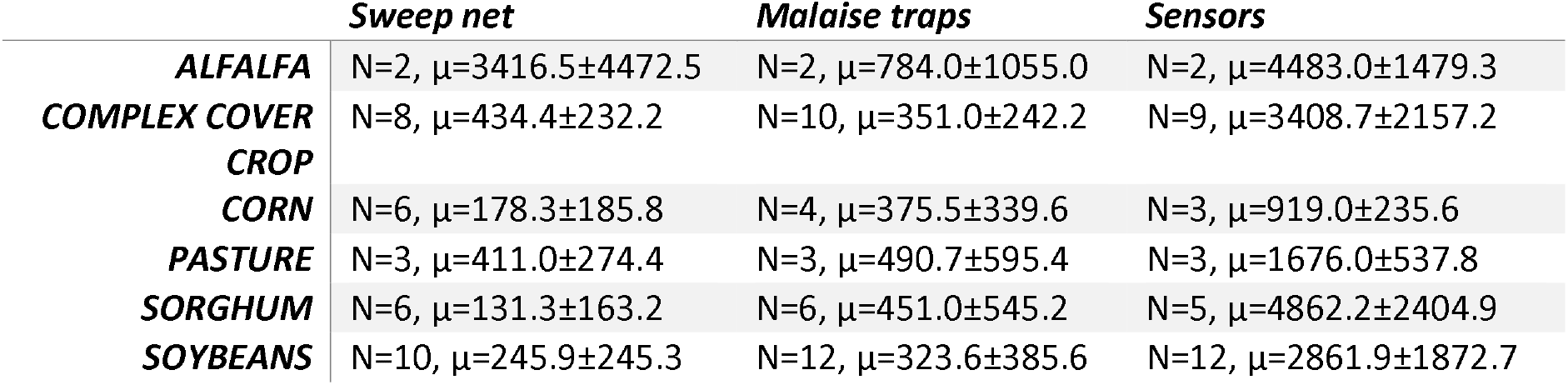
Measured insect abundance per crop and monitoring method. Mean and standard deviation.

**Supplementary Table 3.**
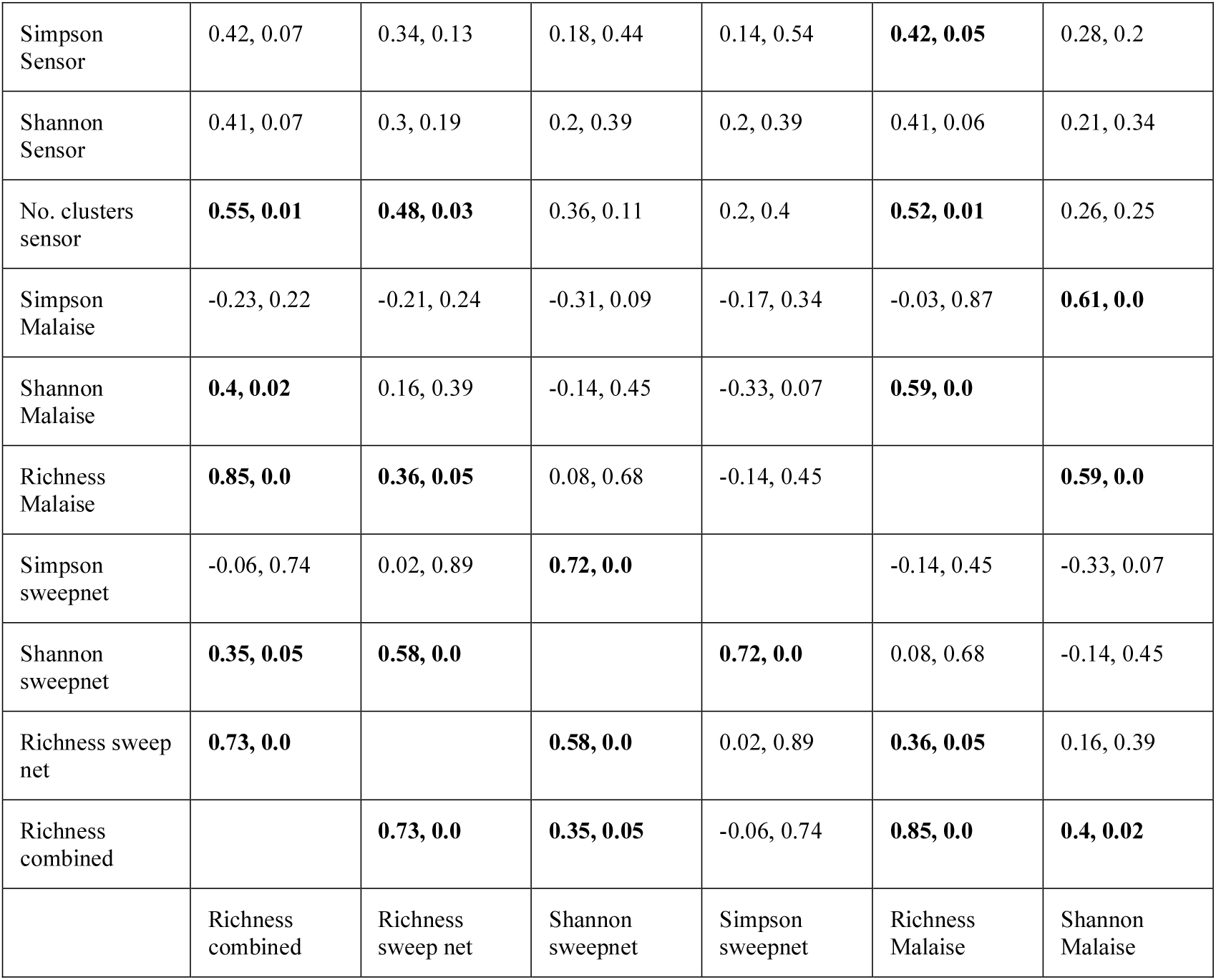
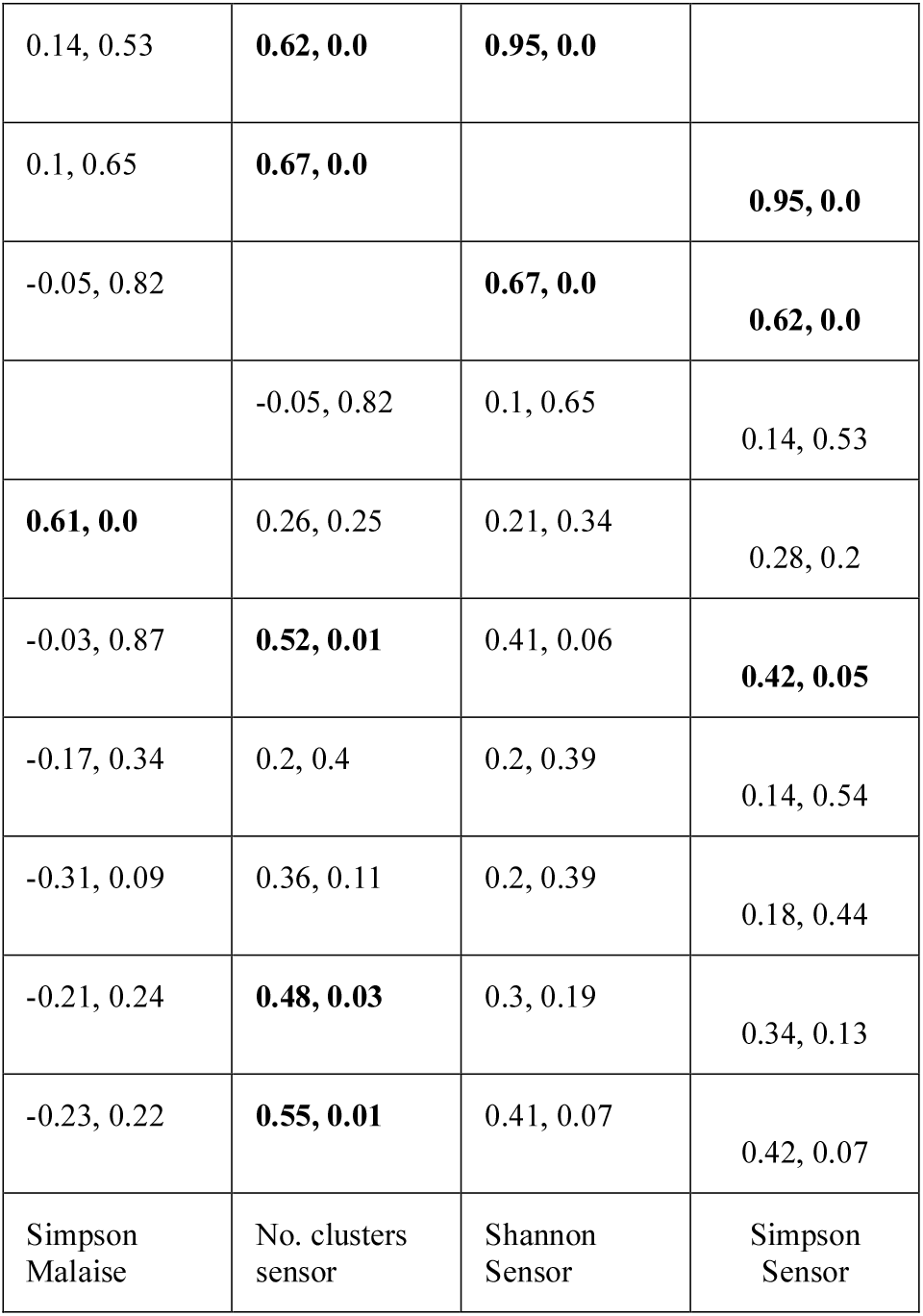
Co-correlations of all biodiversity metrics.

**Supplementary Table 4:** Model parameters for each fitted metric.

| Fitting data | $\epsilon$ | min_samples |
| --- | --- | --- |
| Richness<br>Malaise | 0.007467 | 7 |
| Richness<br>sweep<br>net | 0.009467 | 5 |
| Richness<br>combined | 0.0092 | 5 |
| Shannon<br>Malaise | 0.007467 | 7 |
| Shannon<br>sweep<br>net | 0.0104 | 7 |
| Simpson<br>Malaise | 0.0084 | 3 |
| Simpson<br>sweep<br>net | 5.33E-04 | 3 |

**Supplementary Figure 1.**
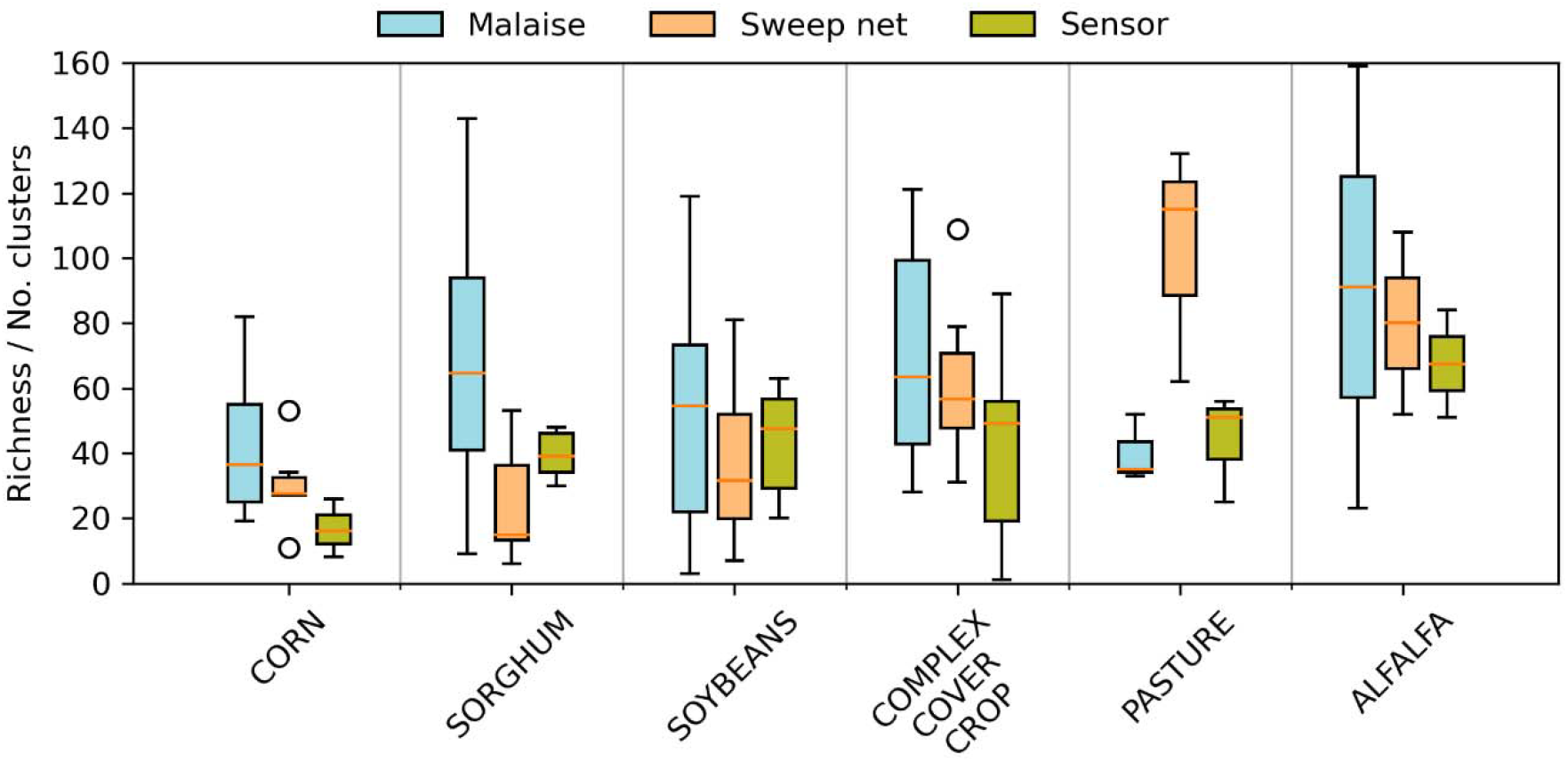
A box plot depicting richness metrics from the Malaise traps, sweep nets, and sensors by field type.

**Supplementary Figure 2.**
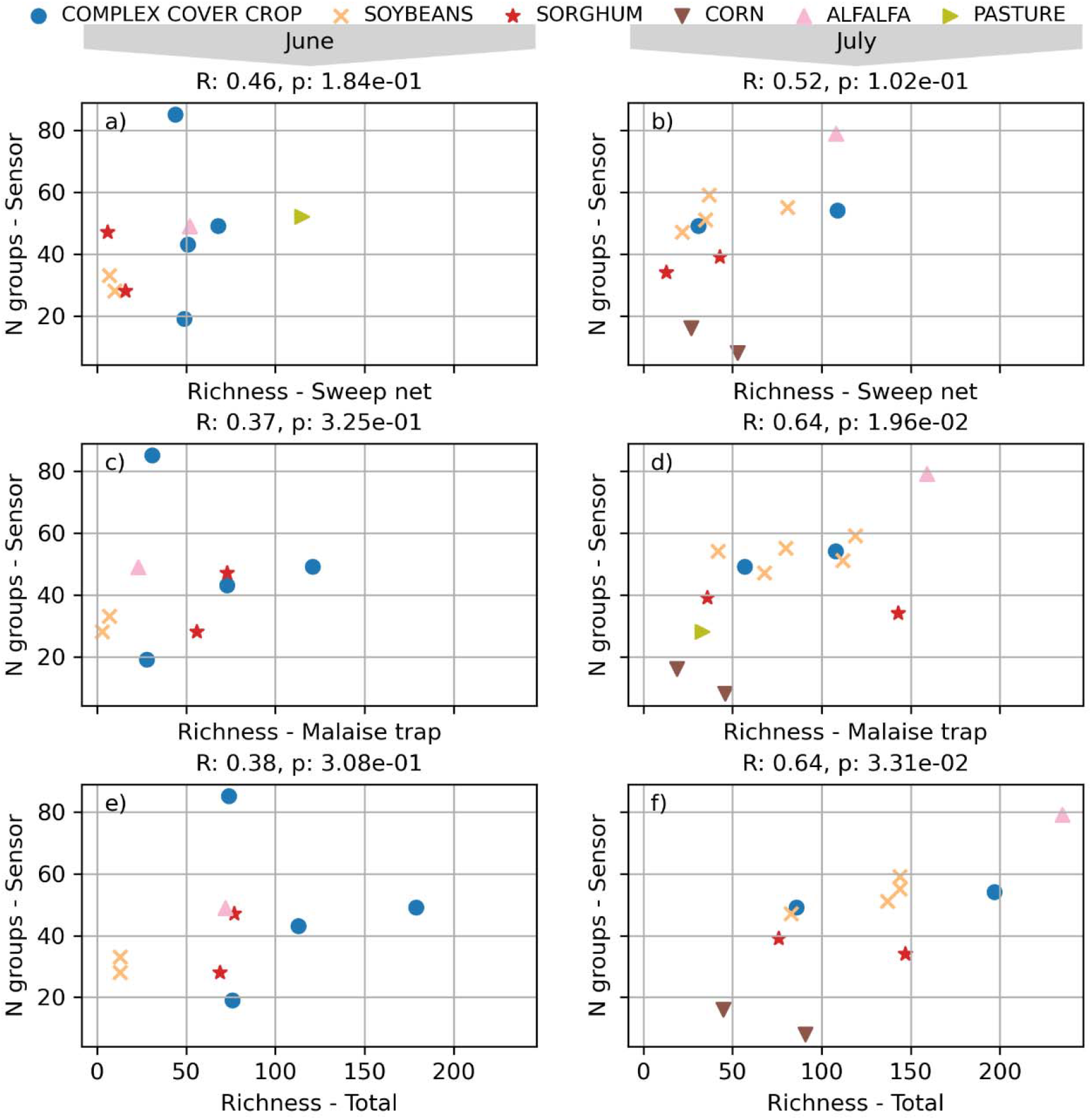
A scatterplot depicting the correlation of the species richness metrics at each field, separated by the June and July timepoints.

